# Diversity of set-related neuronal activity in the dorsal premotor cortex of monkeys during a path-planning task

**DOI:** 10.1101/2024.12.30.630724

**Authors:** Kazuhiro Sakamoto, Naohiro Saito, Hajime Mushiake

## Abstract

Actions involve multiple levels. The dorsal premotor area (PMd) of the primate cerebral cortex is involved in the preparation of actions based on perceptual information. In this study, we recorded neuronal activity from the PMd of monkeys while they performed a path-finding task. In the task, the animals were required to move a cursor step by step to a goal in a maze-like display, and the arm movement and the direction of the cursor operation were dissociated. We observed neurons involved in the preparation of a single arm movement and neurons involved in the preparation of a single cursor operation. Cursor operation-related neurons were also obtained from the lateral prefrontal cortex (lPFC), but the PMd neurons showed more preparatory properties during the execution period than the lPFC neurons. Furthermore, we found neurons involved in the preparation of a specific sequence of cursor movements. The PMd included not only all types of single-shot preparatory neurons but also preparatory neurons for a variety of complex sequences for arm movements and cursor manipulation. These results suggest that the PMd contains diverse neurons involved in different levels of action preparation. Such diversity is one of the critical neural bases for robust goal-directed behavior in complex environments.

**Significance:** There are different levels in actions. The dorsal premotor cortex (PMd) is involved in the preparation of actions based on perceptual information. Here, we recorded neuronal activity from the PM d of monkeys while performing a path-finding task, i.e., moving a cursor step by step to a goal in a maze-like display. In addition to neurons involved in the preparation of a single arm movement and neurons involved in the preparation of a single cursor operation, cells involved in the preparation of a specific sequence of cursor movements were observed. These results suggest that the PMd contains diverse neurons involved in different levels of action preparation. Such diversity is one of the critical neural bases for robust goal-directed behavior in complex environments.

## Introduction

There are different levels of actions. For example, even in the simple case of driving to the nearest supermarket, not only a single action such as turning the steering wheel at an intersection or turning the car to the left at the cognitive level, but also the complex, semantic, and purposive “I’m going to the supermarket” can be considered as a single action. You cannot reach the supermarket without turning the steering wheel and turning the car, but you cannot judge whether it is right to turn left at this intersection without considering the route to the supermarket. Unless you consider actions at different levels, you cannot robustly achieve your goal when errors or changes in the environment occur.

The dorsal premotor area (PMd) of the cerebral cortex is thought to be an important brain region for this. The PMd plays an important role in the preparation and selection of actions by associating arbitrary perceptual information with actions (1–4): PMd-lesioned monkeys are unable to perform conditioned behavior, i.e., to associate visual information with the movement to be performed (5–9); microstimulation of the PMd during behavioral preparation affects the initiation of specific actions (10–15). Correspondingly, many neurons in the PMd show preparatory or set-related activity, i.e., activity that persists after a signal to perform a certain movement is given until the initiation of that movement (16–20). In contrast to cells in the ventral premotor cortex (PMv), which show activity related to visual spatial information and movement dynamics such as force (21,22), these cells are thought to play a role in integrating the necessary sensory and memory information, transforming it into movement and action, and preparing it (23–27). Preparatory activity is elicited not only by a single stimulus that directly indicates the direction of movement, but also by any perceptual cue, such as color (28–30), or by a combination of cues (31–37). This sustained activity does not necessarily reflect the visual nature of the instruction signal, attention to the stimulus, or the upcoming movement (24,38–44). It reflects the intention to perform an action (including the intention of others) (45–51) and the certainty of the action choice (52), and varies with the task (53). For example, in tasks requiring judgments about the size of numbers, even activities corresponding to arithmetic operations are observed (54,55). There are also cells that prepare actions that are not to be performed immediately (24, 56).

Some of these PMd cells encode potential actions, i.e., cells that fire when the possibility arises that a certain behavior must be performed, but stop firing when another cue negates the possibility (57–59). These “potential” cells suggest that the PMd represents possible actions in a parallel and distributed manner (60–63), but there is also the idea that the PMd does not prepare multiple actions, but rather prepares one action and switches between them as needed (64). As the saying goes, “well prepared means no worries”, there is an advantage to preparing possible actions in a parallel and distributed manner in terms of robustness to environmental changes. However, the behavioral task used to record potential cells was simple, in which a cue dictates an action, so there is little need to prepare and select actions at different levels. The idea of preparing a single action and switching it on demand may also have advantages in terms of conserving neural resources.

In this study, PMd cells were recorded while performing a path-planning task (maze task). The task requires the stepwise movement of a cursor to a final goal in a grid display. The movement for cursor manipulation was switched after a certain number of trials to dissociate the arm movement from the cognitive operation of the cursor. In other words, this task allows us to consider actions at different levels (65–70). The analysis revealed not only neurons involved in the preparation of arm movements and the cognitive control of the cursor, but also neurons with complex responses reflecting the preparation to move along a specific path towards the final goal. The properties of these cells are discussed in comparison to lateral prefrontal cortex (lPFC) cells that have connections with the PMd (71–74).

## Results

### Path-Planning Task

Two monkeys (*Macaca fuscata*) were trained to perform a path-planning task that required the planning of multiple cursor movements, controlled using manipulanda, to reach a goal within a maze (Fig. 1A). To begin the trial, the animals were required to hold the two manipulanda in a neutral position for 1 s (initial hold). Subsequently, a cursor was presented at the center of the maze (start display). One second later, the position of goal cursor was presented for 1 s (final goal display). After a delay (delay 1 or 2), the color of the cursor changed from green to yellow, which served as the initiation signal (1st go). After a 1-s hold period, the next go signal was presented (2nd go). When the cursor reached the final goal position, animals received a reward (reward).

**Figure 1.**
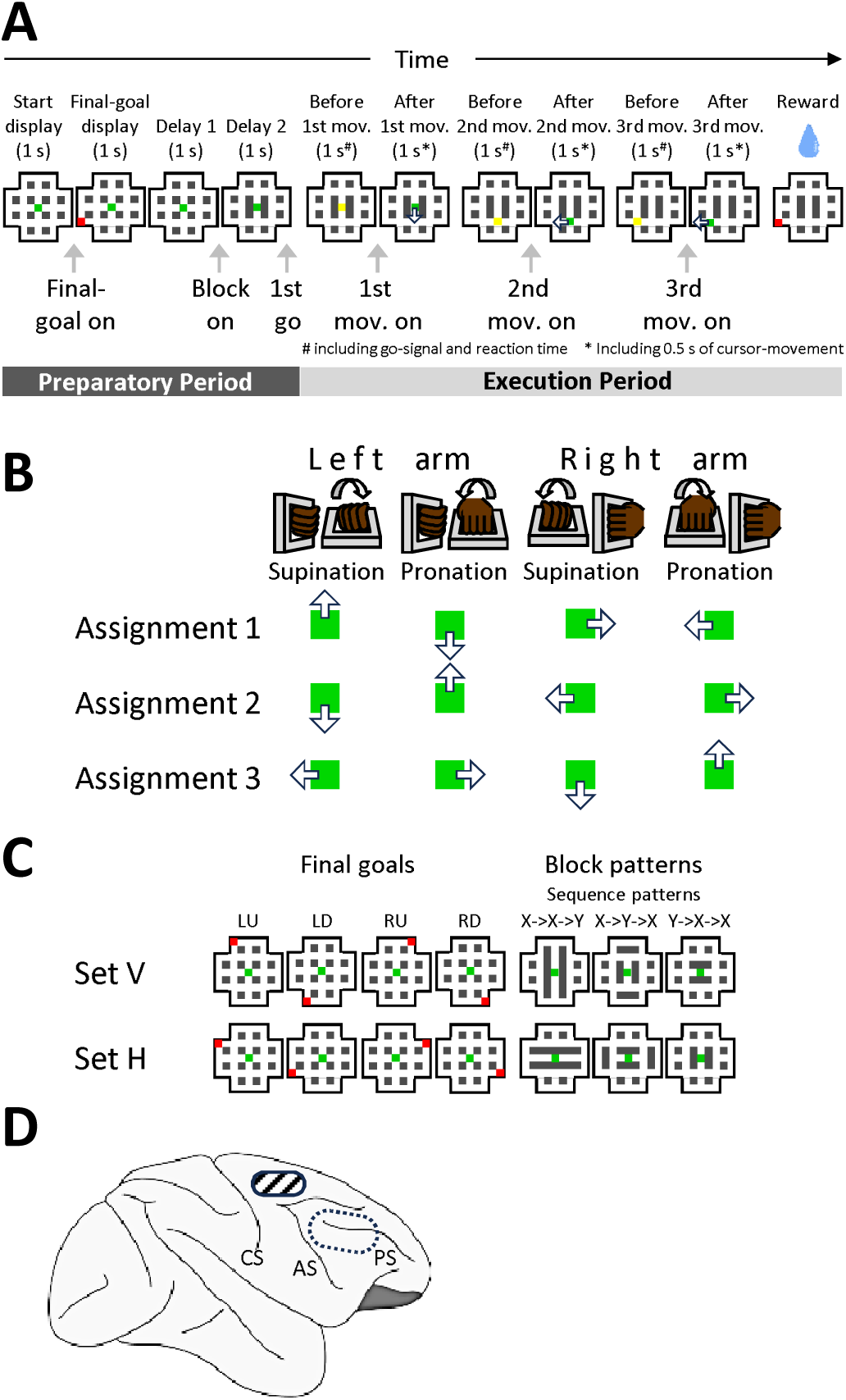
Path-planning task. A. Temporal sequence of events during the task. The behavioral sequence is depicted from the left to the right. Each panel represents a maze that was displayed on a computer monitor. Green and red squares denote the start position and the location of the final goal, respectively. Yellow squares represent a signal for the initiation of movement (GO signal). Empty arrows in the maze indicate the direction of cursor movement. B. Directions of cursor movements (empty arrows) assigned to supination or pronation of either arm. Arm movements were assigned to each cursor movement in three different ways. C. Goals and path blocks used in a recording session. Two sets of display of goal positions (Goals LU, LD, RU and RD) and path blocks (X→X→Y, X→Y→X, Y→X→X). Either set V or set H was used while recording the neuronal activities. Note that the block pattern specifies the sequence pattern of cursor operations. D. Schematic drawing of the right hemisphere. The recording areas of the PMd and the lPFC are indicated by a hatched area and a dotted ellipse, respectively. PS: principal sulcus. AS: arcuate sulcus. CS: central sulcus.

The animal was allowed to make only one of four movements in response to a Go signal: left arm supination (LS), left arm pronation (LP), right arm supination (RS), and right arm pronation (RP). Each movement corresponds to a cursor movement up (U), down (D), left (L), or right (R), and each movement can move the cursor to the next intersection. Arm-cursor assignments were altered after completing a block of 48 trials to dissociate arm and cursor movements. The three assignments were changed in the order 1→2→3→1… as shown in Figure 1B. The data included in this paper were obtained during trials using block patterns that specified the order of actions. Three-block patterns were used in both Set V and Set H (Fig. 1C right). Each pattern specified the shortest sequence as X→X→Y, X→Y→X, and Y→X→X. For example, in the X→X→Y type block of Set V, the shortest cursor operation is U→U→L for the upper left final goal. In→ 89% of trials, monkeys reached the final goal within the minimum number of steps of three.

### Database

Subsequent analyses included neurons from PMd and lPFC that recorded a sufficient number of trials with three arm-cursor assignments (Fig. 1D). Regression analysis (see Materials and Methods) revealed that 334 of 667 recorded PMd cells showed significant changes in firing activity associated with arm movement or cursor manipulation in any of the seven periods (7 s in total, see Fig. 1A) of delay 2, before 1st movement, after 1st movement, before 2nd movement, after 2nd movement, before 3rd movement, and after 3rd movement.) Of these, 12% (83 cells) were related to arm movement only, 23% (153 cells) were related to cursor manipulation only, and 15% (98 cells) were related to both. For comparison, lateral prefrontal cortex (lPFC) cells recorded under the same conditions were also analyzed. Of the 229 cells recorded, 178 lPFC cells changed their activity in response to significant arm movement or cursor manipulation. Of these, about half (50%) (150 cells) were associated with cursor manipulation only, while only 6% (18 cells) and 3% (10 cells) were associated with both arm movements and cursor manipulation or arm movements only, respectively.

### Single-Shot Arm Movement Neuron

Figure 2 shows an example of a cell related to a single arm movement. Figure 2A is sorted for the four arm movements (see figure legend for details). The fluctuation in firing of this cell was certainly not small, but it showed a characteristic and reproducible change when the animal performed a left arm pronation movement (Fig. 2A, lower left, circled in green). This cell increased its activity until just before the start of the left arm pronation movement in response to the Go signal immediately after, and decreased its activity when the movement started. Thus, this neuron is associated with the preparation of a single left arm pronation movement, independent of the step. Reflecting this activity, a regression analysis was performed and the time change in selectivity was plotted as a normalized *F*-value, and the selectivity of the arm movement performed at each step increased, i.e., the selectivity for the first movement increased just before the start of the first movement (Fig. 2B light green), the selectivity for the second movement increased just before the start of the second movement (Fig. 2B green), and the selectivity for the third movement increased just before the start of the third movement (Fig. 2B dark green). These increases in arm selectivity are more pronounced than the increase in cursor operation selectivity (Fig. 2C).

**Figure 2.**
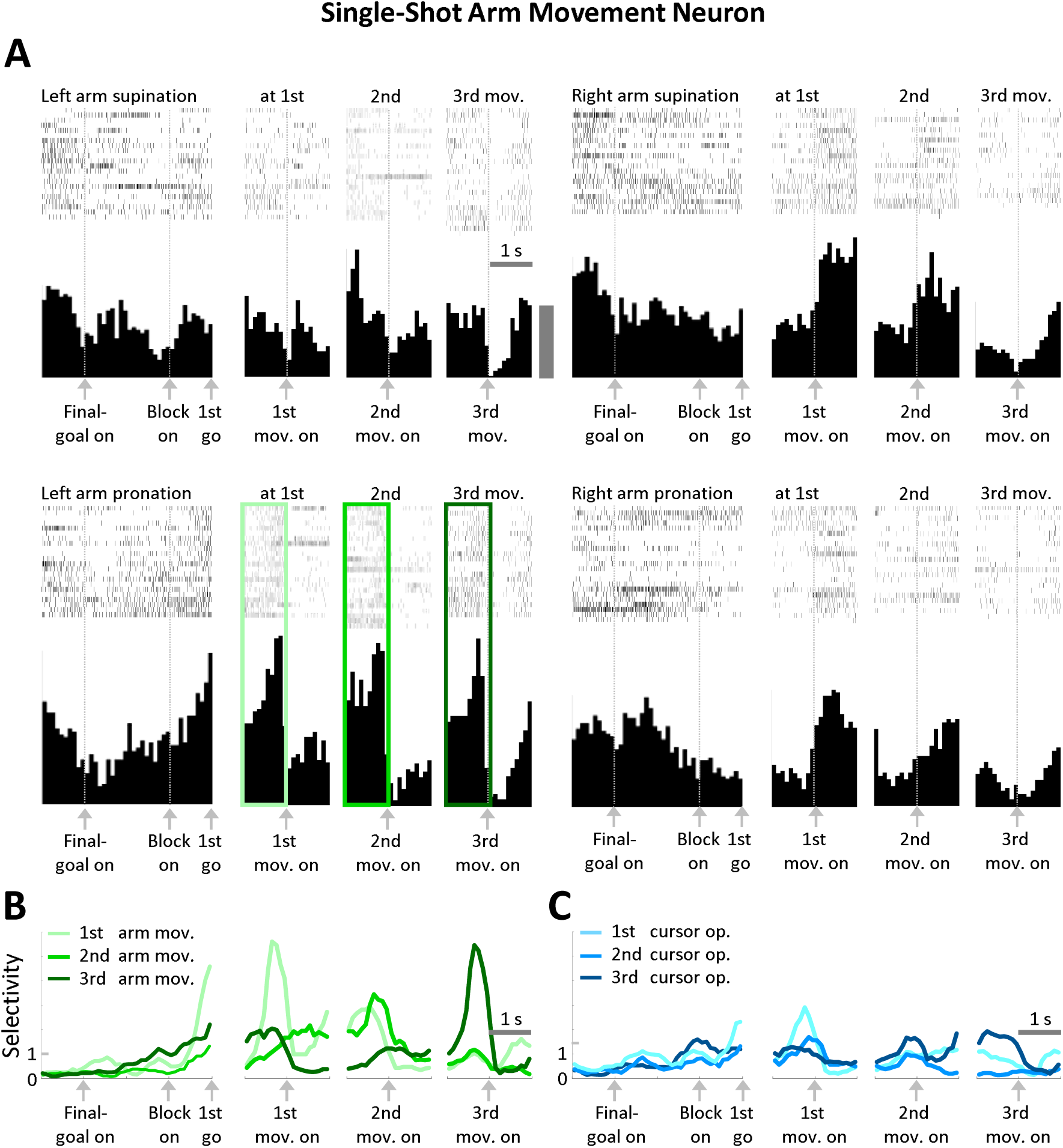
Representative example of a single-shot arm movement neuron. A, Raster plots and spike density histograms of neuronal activity for arm movement at each step. For example, the left-top four panels are of left arm supination, and the panels show trials in which left arm supination was performed in steps before and after each movement onset. Note that the panels are not necessarily from the same trial. However, the raster of the preparatory period is from the same trial as the panel for the first step. This cell showed characteristic activity in the left arm pronation in each step, as indicated by the colored boxes. Gray bar, 10 spikes/s. B. Time course of the 1st (light green), 2nd (green) and 3rd (dark green) arm movement selectivity of the neuron. C. Time course of the 1st (light blue), 2nd (blue) and 3rd (dark blue) cursor operation selectivity of the neuron.

### Single-Shot Cursor Operation Neuron

Figure 3 shows a cell related to a single cursor operation. Figure 3A is sorted for the four cursor manipulations. Similar to the neuron shown in Fig. 2, this cell was highly active until just before the operation to move the cursor to the left in response to the following Go signal, and its activity decreased with the start of the operation (Fig. 3A, lower left, circled in blue). Namely, this cell is involved in the preparation of a single leftward cursor operation, independent of the step. Correspondingly, the regression analysis showed that the selectivity of cursor operations performed at each step increased (Fig. 3B). The selectivity of this cell for arm movements did not increase as much as the selectivity for cursor operations (Fig. 3C).

**Figure 3.**
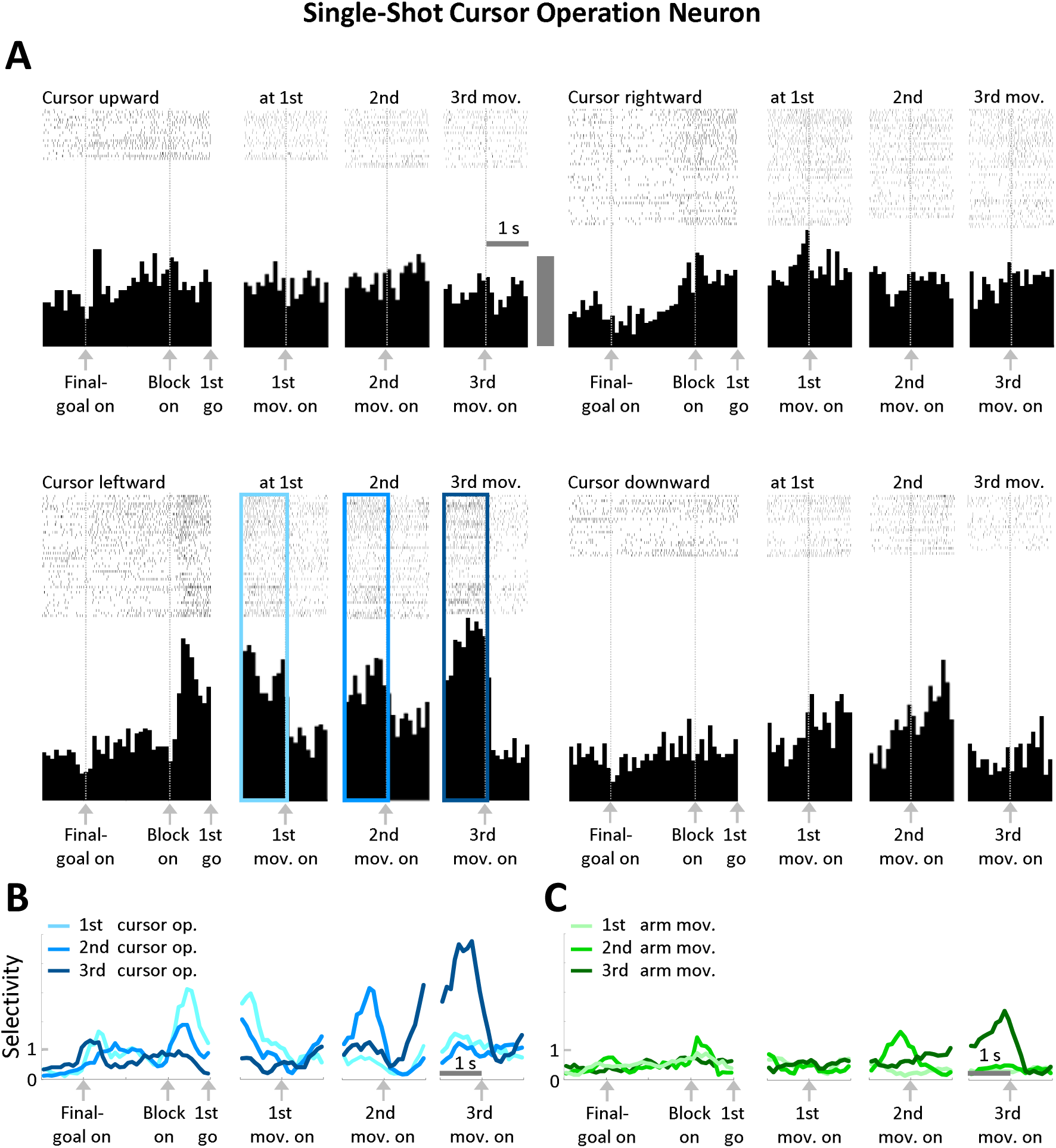
Representative example of a single-shot cursor operation neuron. A, Raster plots and spike density histograms of neuronal activity for arm movement at each step. For example, the left-top four panels are of the upward operation of the cursor, and the trials in which the upward operation was performed in steps before and after each step are collected. However, the raster of the preparatory period is from a trial in which the upward operation was performed in the first step. This neuron showed activity characteristic of leftward operations in each step, as indicated by the colored frames. Gray bar, 5 spikes/s. B. Time course of the 1st (light blue), 2nd (blue) and 3rd (dark blue) cursor-operation selectivity of the neuron. C. Time course of 1st (light green), 2nd (green) and 3rd (dark green) arm movement selectivity of the neuron.

Some cells involved in single-shot movements/manipulations have a step-dependent selectivity. The silent cells shown in Figure 4 showed a marked increase in activity when it was found that the cursor had to be moved down in the first step when a block appeared. The activity gradually decreased until the start of the first movement and was already silent in the second and third steps (Fig. 3A, lower left, circled in blue). Of course, the regression analysis showed that the selectivity of the arm movement was very low (Figure 3B) and that the selectivity of the first cursor operation was very high when the block appeared (Figure 3C, light blue).

**Figure 4.**
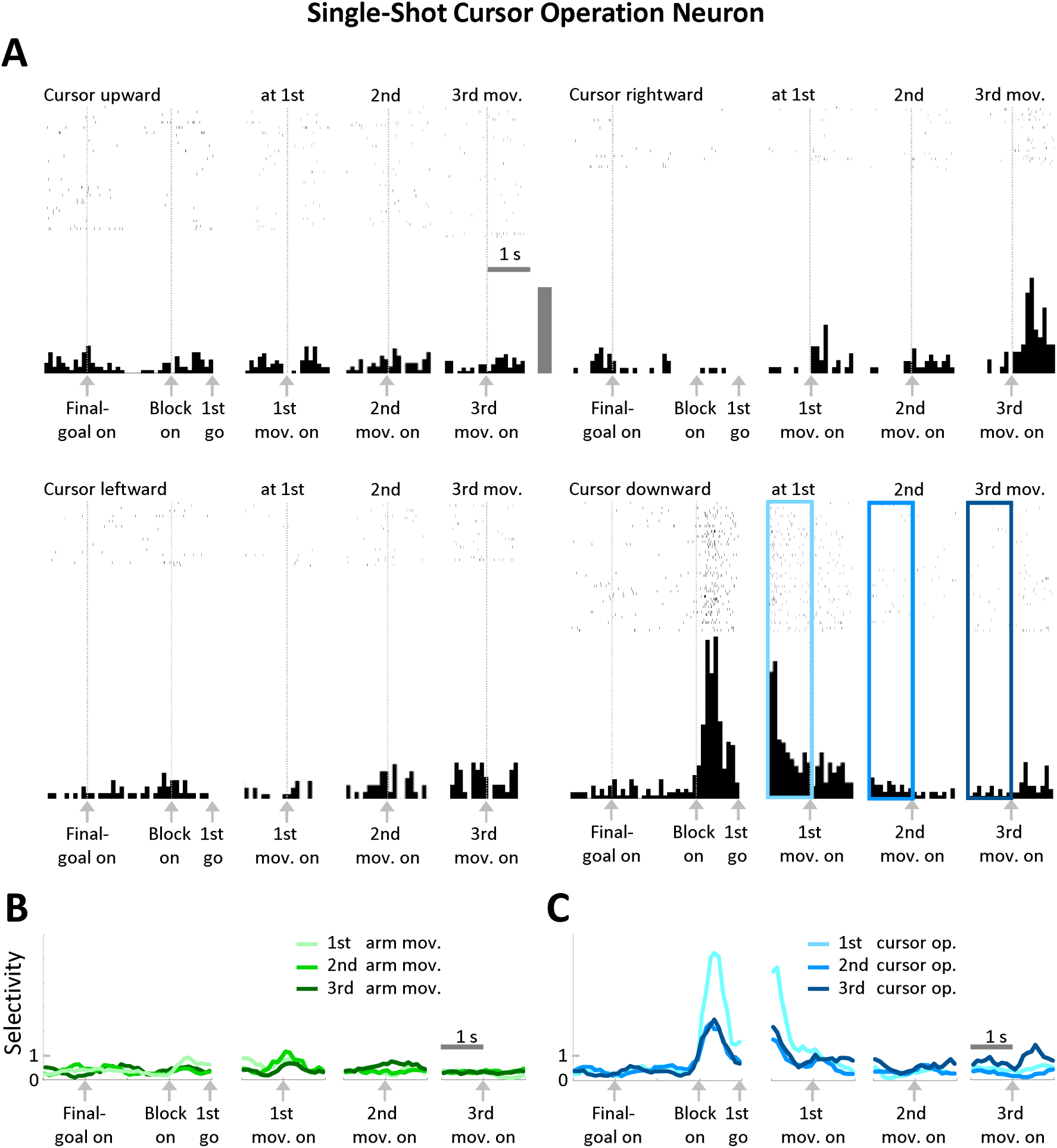
Another example of a single-shot cursor operation neuron. A, Raster plots and spike density histograms. The format is the same as in Fig. 3. This cell showed selective activity only during the first downward operation during the period. Gray bar, 5 spikes/s. B. Time course of the 1st (light green), 2nd (green) and 3rd (dark green) arm-movement selectivity of the neuron. C. Time course of the 1st (light blue), 2nd (blue) and 3rd (dark blue) cursor-operation selectivity of the neuron.

### Comparison of PMd and lPFC in Single-Shot Neurons

Figure 5 shows the time course of selectivity for single-shot cells, i.e., cells that were significantly selective for the i-th arm movement or cursor operation during the before *i* th movement (1 s) or after *i* th movement (1 s) periods.

**Figure 5.**
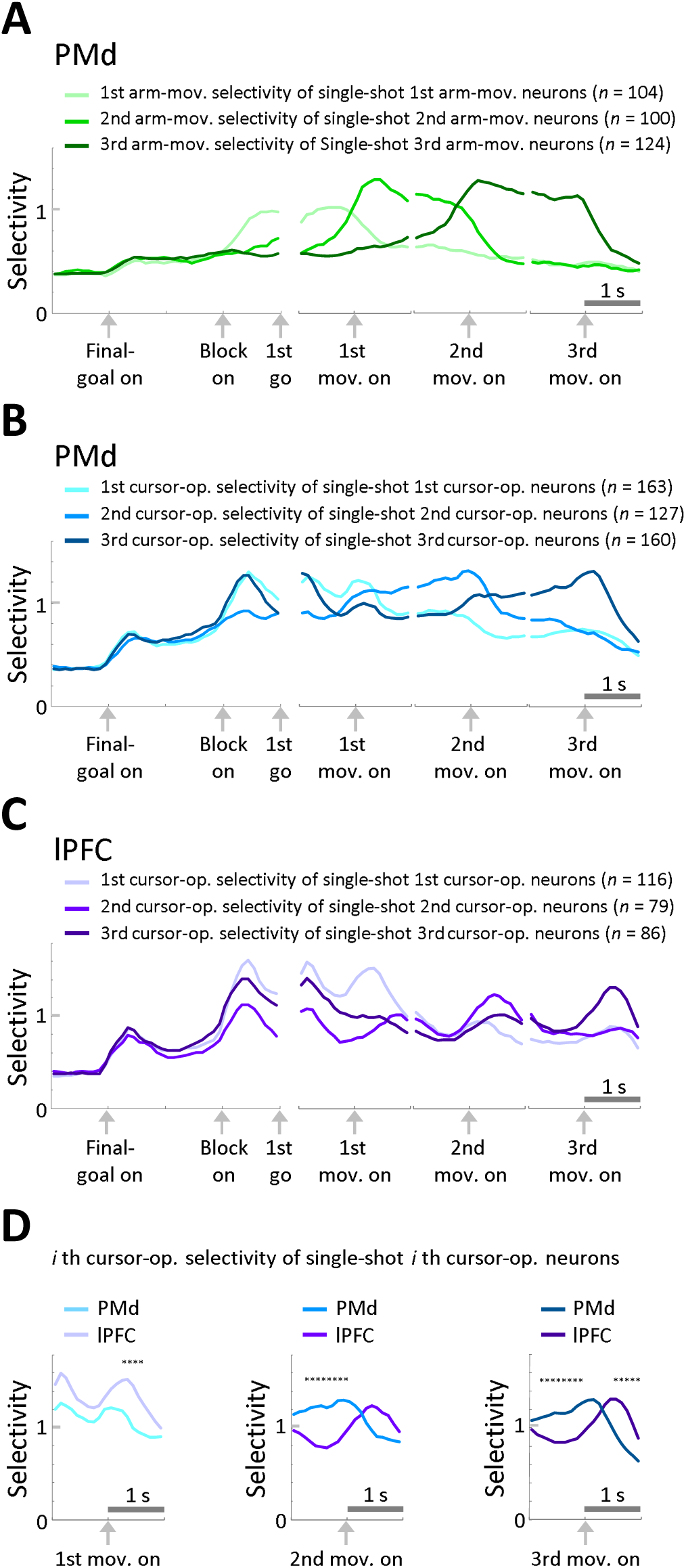
Changes in the arm movement and cursor operation selectivity of single-shot neurons at the population level. A, Time course of the i-th arm movement selectivity of the single-shot i-th arm movement neurons of PMd. B. Time course of the *i*-th cursor operation selectivity of the single-shot *i*-th cursor operation neurons of PMd. C. The same plot for lPFC neurons. D. Comparison of cursor operation selectivity at each step between PMd and lPFC. Times that were significantly different (p < 0.05) are marked with an asterisk.

Figure 5A shows the change in selectivity of the single-shot arm movement neurons. First movement selectivity increased with the presentation of the block and decreased to baseline at the onset of the first movement. Second movement selectivity increased at the onset of the first movement and decreased at the onset of the second movement. Similarly, third movement selectivity increased with the onset of the second movement and decreased with the onset of the third movement. All arm movement selectivities showed a “preparatory” profile, in which selectivity increased with a significant event, such as the presentation of a block or the onset of the previous movement, then remained high, and then decreased with the execution of the movement of interest.

Figure 5B is of the single-shot cursor operation neuron. Although the selectivity for the first movement was somewhat transient, it had a “preparatory” nature similar to that of the arm movement preparatory neuron above. That is, the selectivity increased near the start of the previous operation, remained high, and then decreased with the start of the cursor operation.

In contrast to arm movement selectivity, cursor operation selectivity is also prominent in the lPFC (67). Figure 5C shows the time course of selectivity for single-shot cursor operation neurons in the lPFC. Selectivity for each hand first increased transiently after block presentation, but then decreased once. While the first movement is difficult to distinguish because it is affected by the block presentation, the selectivity for the second and third movements does not appear to maintain high selectivity from the onset of the previous operation; rather, selectivity appears to increase after the onset of that movement.

To make this point easier to understand, in Fig. 5D, Figure 5D compares the cursor operation selectivity of PMd and lPFC at each step. As can be seen in the figure, the selectivity of lPFC exceeds that of PMd after the start of the operation. The selectivity of PMd is significantly higher for the second and third movements, which are not affected by the block presentation (times where p < 0.05 in the *t*-test are indicated by *). These results suggest that PMd is more “preparatory”, at least in the execution period, i.e., in the period after the 1st Go.

### Complex Sequence Neuron

However, not all cells observed in PMd are selective for only a single action as described above. The example shown in Fig. 6 is a neuron that is selective for down→left→left cursor movements, that is, that responds selectively to a particular cursor operation sequence. As shown in Fig. 6A, when the final goal was presented at the LD, this cell started to fire because there was a possibility of performing a down→left→left cursor sequence. However, when a block was presented and it was still possible to perform a down→left→left operation, the cell continued to be active, and finally, when the third movement was started, its activity rapidly decreased. On the other hand, when a block was presented and it turned out that the down→left→left operation could not be performed, it rapidly decreased its activity. In this sense, this cell encodes a “potential” cursor operation sequence. Plotting the change in cursor operation selectivity for each step, the selectivity for the first step remained high until the third step (Fig. 6B). This is because it is the first cursor operation that distinguishes the down→left→left operation from the other two sequences (however, the reason why the selectivity for the third cursor operation was high just before the onset of the first movement is that the decrease in firing at that time was not sufficient for the left→down→left pattern, which deviated from the impression one gets from the PSTH, and rather the large decrease in activity for the left→left→down pattern in the first step distinguished it from the other two patterns). Note that in the following, cells involved in complex operations are defined as cells that show selectivity for the i-th operation other than the i-th step. Figure 6C shows the results of a new regression analysis of final goal selectivity and cursor sequence down→left→left selectivity for this neuron. When the LD goal was presented and this neuron had the possibility to perform down→left→left until the block was presented, the cell fired well in all cases, resulting in the high selectivity for the final goal(Fig. 6C red). However, when the block was presented, the possibilities of left→down→left and left→left→down were eliminated, so the high selectivity for the final goal was replaced by high selectivity for down→left→left (Fig. 6C purple).

**Figure 6.**
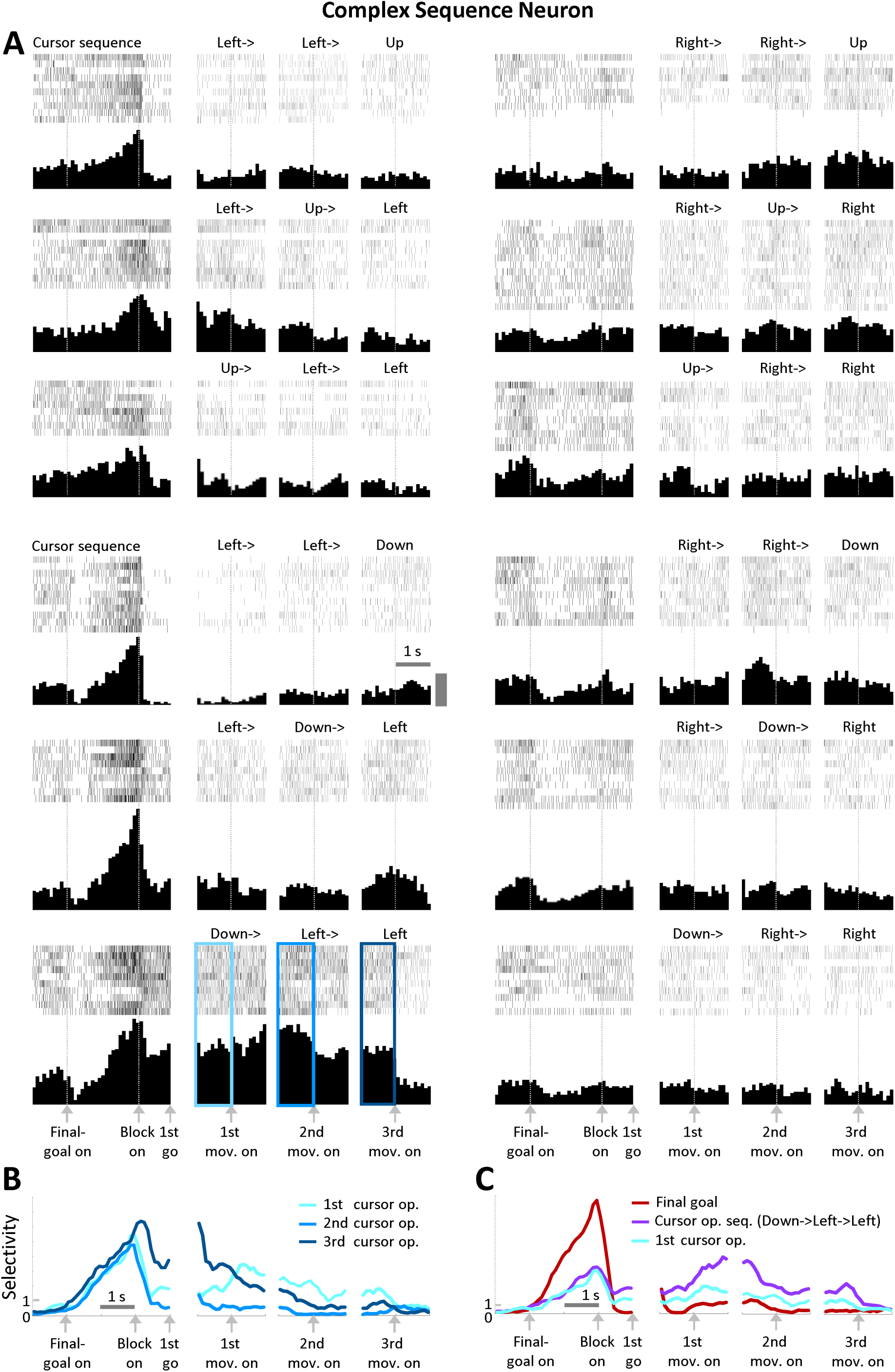
Representative example of a complex sequence neuron. A, Raster plots and spike density histograms of neuronal activity. Panels are sorted according to each sequence of cursor operations. Note that unlike Figs. 2-4, the four panels in a row are from the same trial. This cell showed persistent activity for the down→left→left sequence, as indicated by the colored frames. Gray bar, 10 spikes/s. B. Time course of the 1st (light blue), 2nd (blue) and 3rd (dark blue) cursor operation selectivity of the neuron. C. Time course of the neuron’s final goal (red), cursor-operation sequence of down→left→left (purple) and 1st cursor-operation (light blue, same as in B) selectivities.

### Numbers of preparatory patterns and preparatory neurons

The number of patterns of preparatory activity and the number of neurons that showed them were counted. Patterns of preparatory activity were assigned to each cell by determining whether the activity was selective for each of the three periods: before the 1st movement, before the 2nd movement, and before the 3rd movement. For example, the arm movement selectivity of the single-shot arm movement in Figure 2 was set to 123 because its activity showed significant changes in arm movement for each step. The cursor movement selectivity of the single-shot cursor movement cell in Fig. 4, which was significant only for the first step, was set to 100, and the cursor operation selectivity of the complex sequence neuron in Fig. 6 was set to 311. There were a total of 4^3^ – 1, or 63, patterns of arm movement and cursor operation. Of these, single-shot patterns were defined as patterns that showed selectivity for the current step and no selectivity for the other steps, and specifically included the seven patterns 100, 103, 120, 123, 20, 23, and 3. The remaining 56 patterns were included in the complex sequence patterns. The PMd neurons had all seven single-shot patterns for both arm movements and cursor operations. 57 single-shot preparatory cells for arm movement and 75 single-shot preparatory neurons for cursor manipulation were included. On the other hand, the PMd showed 27 (71 neurons) and 38 (99 neurons) complex sequence patterns for arm movement and cursor manipulation, respectively, accounting for 48% and 68% of the 56 possible patterns.

## Discussion

In this study, we recorded neuronal activity in the PMd of monkeys during a path-planning task. Analysis revealed a number of cells suggesting that the PMd is involved in multiple action preparations. Specifically, we observed not only set-related activity corresponding to single arm movements and cursor operations, but also a large number and variety of set-related cells corresponding to complex sequences. Cursor operation-related neurons were also observed in the lPFC, but PMd cells showed a more “preparatory” profile.

In our previous work and in this paper, we showed that lPFC cells are active prior to execution, especially immediately after block presentation, reflecting individual cursor operations (67). In general, the lPFC is thought to play a crucial role in executive function (75–77). Specifically, the lPFC is involved in the formulation of action plans as the brain’s “executive” (66–70,78,79). In our path planning task, the lPFC shows neuronal activity related to the cursor operation plan for the first to third steps during the block presentation period (delay 2), but is not directly involved in arm movements (66,67). On the other hand, as shown in Fig. 5D, the increase in cursor operation selectivity during the execution period occurred after movement onset, as if the lPFC was confirming that the “subordinate” had actually executed the action.

On the other hand, PMd cells are active at different levels, from cursor operation to arm movement, from single actions to complex actions, from initiation to completion of the action. As shown in Fig. 6, even a complex cursor-operation sequence activates the neuron when it prepares the sequence as a “potential” action, as argued by Cisek et al.(58, 60). This is as if the “capable middle manager” between the lPFC and the primary motor area is ready to respond quickly and in multiple ways to any coarse plans provided by the brain’s “executive". The close anatomical connections between the lPFC and PMd (71–74) also support the idea that they play exquisitely complementary roles in carrying out planned actions, like the executive and the middle manager in a company.

In recent years, as in other cortical areas, there has been increasing discussion of coding by the dynamics of neuronal populations in PMd (80–84). Even if the individual cells that make up a population do not appear to encode much information at first glance, the population dynamics reflect cognitive information, decision making, and even information that predicts subsequent reaction times, *etc*. The decoding of population dynamics itself, which attempts to find significance in seemingly meaningless neurons and extract rich information, is significant and useful when considering the use of PMd neural activity for brain-machine interfaces (85,86). However, population coding can only decode predetermined dimensions (such as task parameters). What the authors, as neurophysiologists, want to explore is the brain’s way of information processing that underlies its adaptability to unpredictably changing environments.

There is a Chinese proverb that says, “Even the wisest man has one mistake in a thousand thoughts. No matter how talented a person is, he will make mistakes. It is expected that the wisdom to avoid such mistakes and adapt to major changes in the environment will be generated, which is why democracy and diversity are loudly called for in modern society. Even a group of bees born from the same queen has diversity and democracy to adapt to their environment (87). When the bees move from hive to hive due to environmental changes, the worker bees go exploring in different directions beforehand, and each of them brings information to form a “consensus” through dance, and the whole group moves accordingly. What the neural population dynamics analysis sees is similar to the behavior of a bee population. However, it should be noted that the reason why their decision is accurate in a complex environment is due to the fact that worker bees bring back information from a variety of directions. The essence of their environmental adaptability lies not in the population dynamics, but in the diversity of the worker bees’ explorations. The diversity of set-related neurons observed in this study shows an important aspect of the high environmental adaptability of the brain. The results of this study suggest that a variety of decision-making and behavioral selection mechanisms exist within a single individual. Artificial intelligence (AI) is currently developing rapidly and becoming widespread in society. However, it can also make surprisingly rash and incorrect decisions. We believe that the results of this study will have important significance in helping AI acquire the ability to make more reliable decisions in the future. Indeed, no matter how complex the path-planning task was, the monkeys have been overtrained, so the task may not have been so uncertain for them. However, the multiplicity of preparation and diversity of preparatory neurons shown in this study are thought to be one of the foundations of good intracerebral deliberation (48) and high environmental adaptability.

## Acknowledgements

This work was supported by JSPS KAKENHI Grant Number 20K07726, 23K06001 (Kiban C) 23K18159 (Grant-in-Aid for Challenging Research (Exploratory)), MEXT KAKENHI Grant Number 20H05478, 22H04780 (Hyper–Adaptability) and SIP Project Phase 3 (Development of Quantum Spin Sensor and Development and Demonstration of Use Cases).

## Materials and Methods

### Subjects and experimental setup

This study was performed on two male Japanese monkeys, *Macaca fuscata*, 6–7 kg. During the experiment, each monkey was seated in a primate chair with its head restrained and was oriented toward a computer monitor on which a checkerboard-like maze and a cursor were displayed. Each monkey could operate two manipulanda in the chair by the supination and pronation of either forearm with one degree of freedom to move a cursor through the maze. The supination and pronation of each forearm were assigned to four cursor directions, and eye position was monitored using an infrared eye camera (R21-C-AC; RMS) with a 250 Hz sampling rate. All experimental protocols were approved by the Animal Care and Use Committee of Tohoku University (Permit #20MeA-2), and all animal protocols conformed with the National Institutes of Health guidelines for the care and use of laboratory animals, as well as with the recommendations of the Weatherall Report.

### Recordings and Data Analysis

We used conventional electrophysiological techniques to obtain in vivo single-cell recordings from the PMd and lPFC regions of the right hemisphere described elsewhere (66)(Figure 1D). At each recording site, visual and somatosensory receptive fields were checked, and muscle activity and eye movements were confirmed by intracortical micro-stimulation. These were compared with the sulcus location confirmed by MRI, surgery, and each experimental session to ensure that the recordings were not from surrounding areas such as the frontal eye field, PMv, or primary motor area.

Multiple regression analysis was performed on spike counts for each 100-ms time window to assess the relevance of neuronal activity with arm movement or cursor manipulation. The following 12 analyses were performed: first arm movements, combinations of final goal position and first arm movements, second arm movements, combinations of final goal position and second arm movements, third arm movements, combinations of final goal position and third arm movements, first cursor operations, combinations of final goal position and first cursor operation, second cursor operation, final goal position and second cursor operation, third cursor operation, final goal position and third cursor operation. Neurons that showed an F value of p < 0.05 over three time windows in any of the 1-second intervals of delay 2, before 1st movement, before 1st movement, before 2nd movement, after 2nd movement, before 3rd movement, and after 3rd movement (Fig. 1A) were included in the analysis. We defined “selectivity” as the *F* value obtained, normalized by the *F* value at p = 0.05. All of the selectivity figures are smoothed with a sliding window of 500 ms.

